# Fungal traits important for soil aggregation

**DOI:** 10.1101/732628

**Authors:** Anika Lehmann, Weishuang Zheng, Masahiro Ryo, Katharina Soutschek, Rebecca Rongstock, Stefanie Maaß, Matthias C. Rillig

**Author notes:** ***Author to whom correspondence should be sent*** Anika Lehmann, Freie Universität Berlin, Institut für Biologie, Altensteinstr. 6, D-14195 Berlin, Germany.

## Abstract

Soil health and sustainability is essential for ecosystem functioning and human well-being. Soil structure, the complex arrangement of soil into aggregates and pore spaces, is a key feature of soils under the influence of soil life. Soil biota, and among them filamentous saprobic fungi, have well-documented effects on soil aggregation. However, it is unclear what fungal properties, or traits, contribute to the overall positive effect on soil aggregation. So far, we lack a systematic investigation of a broad suite of fungal species for their trait expression and the relation of these traits to their soil aggregation capability.

Here, we apply a trait-based approach to a set of 15 traits measured under standardized conditions on 31 fungal strains including Ascomycota, Basidiomycota and Mucoromycota, all isolated from the same soil.

We found a spectrum of soil aggregate formation capability ranging from neutral to positive and large differences in trait expression among strains. We identified biomass density (positive effects), leucine aminopeptidase activity (negative effects) and phylogeny as important modulators of fungal aggregate formation capability. Our results point to a typical suite of traits characterizing fungi that are good soil aggregators; this could inform screening for fungi to be used in biotechnological applications, and illustrates the power of employing a trait-based approach to unravel biological mechanisms of soil aggregation, which could now be extended to other organism groups.

## 1. Introduction

Soil is our most vital resource, with soil and its biodiversity contributing to many ecosystem processes (Bardgett and van der Putten, 2014), and to human nutrition, health and wellbeing (Wall et al., 2015). Soil has been described as the most complex biomaterial on Earth (Young and Crawford, 2004) with soil structure as one of its most important features. Soil structure represents the three-dimensional arrangement of soil particles into aggregates and associated pore spaces and is also a crucial parameter for sustainable management of soils (Bronick and Lal, 2005); therefore, it is of great interest to unravel how soil biota contribute to the process of soil aggregation.

Many soil biota influence soil aggregation (Lehmann et al., 2017b), and among them are the filamentous fungi. These fungi have a particularly well-documented impact on soil structure especially at the macroaggregate (>250µm) scale, as highlighted in a meta-analysis (Lehmann et al., 2017b). Soil aggregating capability of fungi is hypothesized to be due to a range of physical, morphological, chemical and biotic traits (Six et al., 2004; Bronick and Lal, 2005; Lehmann et al., 2017a). While foraging and growing through soil, fungi are thought to entangle and enmesh soil particles and aggregates due to their filamentous growth form (Tisdall and Oades, 1982). Fungi also exude extracellular biopolymers which can act as cements and surface sealants for soil aggregates (Chenu, 1989; Caesar-TonThat and Cochran, 2000; Daynes et al., 2012), and enzymes degrading organic matter (Baldrian et al., 2011), which may serve as aggregate-disintegrating agents. Among the molecules they release are also hydrophobins, which can modify wettability of aggregates, likely serving a stabilizing function (Zheng et al., 2016). While growing through soil, fungi also interact with other members of the soil community, for example they can be grazed upon by Collembola, which can also influence soil aggregation ability (e.g. (Siddiky et al., 2012a; Siddiky et al., 2012b)).

Fungi likely differ in many of these traits, and thus also in their soil aggregation capability. In fact, exploring a global dataset of fungal contributions to soil aggregation, Lehmann et al. (Lehmann et al., 2017b) revealed a wide range in soil aggregation effectiveness for the 117 species for which experimental data were available. However, in this analysis it remained unclear which fungal traits underpin the observed effects on soil aggregation, simply because the relevant trait data are unavailable.

What is needed are studies that systematically compare fungal traits in a set of species and that relate these to soil aggregate ability. So far, only a limited number of such studies are available (Table S1). These studies have mainly focused on fungal biomass and some chemical traits, using specific fungal groups, such as arbuscular or ectomycorrhizal fungi. Much less is known for soil saprobic fungi. In all these studies a limited set of fungi (typically in the range of 3 to 9 species) was examined for their traits (no more than 3 traits). In cases where larger suites of fungi (up to 85 fungal strains/ mutants) were investigated for their soil aggregation ability no traits were measured (Table S1). This lack of data currently prevents us from arriving at more broadly generalizable conclusions.

A way forward to address this issue is by applying a trait-based approach, especially for saprobic fungi (Lehmann and Rillig, 2015). As opposed to arbuscular mycorrhizal fungi, for which most work in this context has been done (Rillig et al., 2015), there are also clear traits for disaggregation ability in this group: aspects of enzymatic ability. In a trait-based approach, using a reasonably large suite of isolates, organismal traits can be related to specific functions. Such approaches generally convert species into points in ‘trait-space’, thus overcoming limitations associated with examining a few, idiosyncratically selected strains, and thus allowing for more generalizable inferences (Crowther et al., 2014; Aguilar-Trigueros et al., 2015).

Here, we investigated a set of 31 filamentous fungal strains, all saprobic fungi isolated from the same soil and then compared under identical conditions in the laboratory. The 31 strains are distributed among the Ascomycota, Basidiomycota and Mucoromycota (Spatafora et al., 2016), and we screened each for the expression of a suite of 15 traits. With these data, we wished to determine (i) which morphological, chemical and biotic traits are most important for soil aggregation and (ii) what characterizes an efficient or poor soil aggregator.

## 2. Materials and Methods

### 2.1. Soil and fungal strains

Soil samples and fungal strains were obtained from Mallnow Lebus, a dry grassland in a natural reserve (Brandenburg, Germany, 52° 27.778’ N, 14° 29.349’ E) characterized by a sandy loam soil texture. The collected soil samples were either used for establishing fungal cultures or were air-dried and stored until further use in experiments. The isolation of the 31 fungal strains was previously described in Andrade-Linares et al. (Andrade-Linares et al., 2016). Briefly, washed and diluted soil was used for the isolation procedure to minimize spore abundance and to increase the probability of capturing fungi derived from hyphae attached to soil particles (Gams and Domsch, 1967; Thorn et al., 1996). Afterwards soil suspensions were incubated on a variety of media with applications of different antibiotics suitable for cultivation of Ascomycota, Basidiomycota and Mucoromycota while suppressing bacterial growth. Isolates were grown on PDA at room temperature (22°C). Our final set of fungal strains comprised 20 Ascomycota, four Basidiomycota and seven Mucoromycota strains (Fig.1, Table S2). The corresponding phylogenetic tree was calculated following the procedure by Andrade-Linares et al. (2016). Briefly, ITS regions were sequenced using the primers ITS1F and ITS4. Sequences were matched in GenBank and aligned via Muscle v. 3.8.31 (Edgar, 2004). For reconstruction of phylogenetic relationships across the 31 fungal strains, a Bayesian maximum likelihood approach was applied using BEAST v. 1.7.2 (Drummond and Rambaut, 2007). A general time reversible substitution model was run with gamma-distributed substitution rates. Further a Bayesian chain with 20 million generations was implemented. The phylogenetic tree was rooted by the isolate Chytridium olla (GenBank accession number: FJ822974) which was used as an outgroup. Generated trees were sampled every 2000 generations from which the first 1000 were discarded as the burn-in (see e.g. Nascimento et al., 2017). The summary tree represents the maximum clade credibility tree with median clade heights.

**Fig.1.**
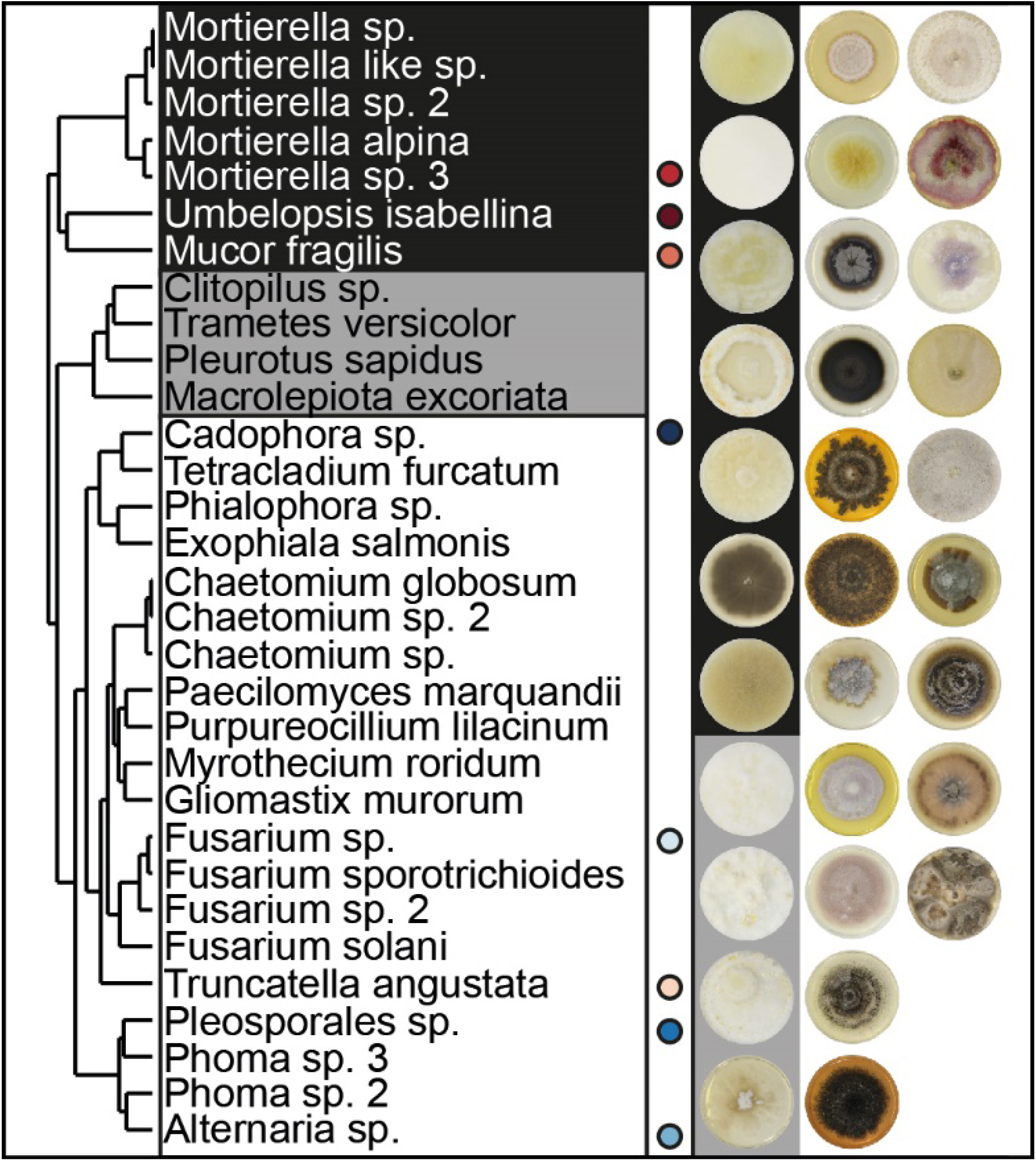
Overview of fungal strains. Phylogenetic tree (maximum clade-credibility tree) of the 31 saprobic fungal strains comprising members of the phyla Ascomycota, Basidiomycota and Mucoromycota. Following the order of the tree, images of four week old colonies grown on PDA are assigned to the tree. Further information about phylogeny and accession numbers of the 31 fungal strains are available in Table S2. Strains performing best and poorest are marked; blue symbols represent good and red symbols poor aggregators.

### 2.2. Soil aggregate formation

The soil aggregate formation assay used here aimed to test for *de novo* aggregate formation by fungi. This technique was modified from Tisdall et al. (2012). Here, we filled 6 cm petri dishes with a 5 mm layer of agar (1.5%, Panreac AppliChem, Darmstadt, Germany) to provide moisture, and this layer was covered with 10.0 g of soil. The soil was gently poured onto the agar to avoid any artificial compaction. Prior to this, the soil (from the field site from which the fungi were originally isolated) was sieved to a fraction < 1mm and autoclaved two times in a dry cycle. The soil was then allowed to equilibrate for two days on the agar before inoculation. During this time, the soil was rewetted by capillary action. This way, we provided a moist but not waterlogged environment for the fungal strains. The fungal strains used for inoculation were cultured on PDA and incubated with sterilized poppy (*Papaver somniferum*) seeds as carrier material. Colonized poppy seeds were transferred to soil - with two seeds per species added per soil plate. For the controls, non-colonized poppy seeds incubated on PDA were transferred to the soil plates. Finally, plates were sealed and stored at room temperature (22°C, the culturing temperature of our fungal strains) in the dark for six weeks until harvest. The experiment consisted of ten replicates for 31 fungal strains and a control, resulting in 320 experimental units.

We visually confirmed for every strain (on two replicates) that hyphae were not just growing on the surface of the soil, but that that mycelium was present inside the soil. At harvest, the plates were opened and dried at 60°C overnight. Subsequently, the soil was carefully extracted from the Petri dishes, passed through a 1 mm sieve to extract all aggregates larger than 1 mm, which were formed during the experiment. To do so, we vertically moved the sieve two times to allow separation while avoiding abrasion of soil aggregates. Additionally, we tapped against the sieve frame. By this, we increased the likelihood of passing aggregates and particles <1mm captured by hyphae through the mesh. The weight of the soil fraction >1mm was used for the calculation of the soil aggregate formation for our 31 fungal strains and the corresponding controls following the equation: % SAF = (aggregates _>1mm_/10.0) *100.

This approach offers the opportunity to test soil aggregate formation for an *a priori* size fraction (here 1mm). However, this design does not capture any dynamics for the <1mm soil fraction. Hence any impact of the 31 fungal strains on e.g. microaggregate formation could not be evaluated here.

### 2.3. Trait measurements

To build a trait database, we investigated 15 different traits capturing morphological, chemical and biotic features of our 31 fungal strains (Lehmann and Rillig, 2015; Lehmann et al., 2017a). The traits were chosen to characterize different aspects of the fungal mycelium and its products by which the fungus interacts with its environment. Additionally, the traits had to be measurable for all 31 strains, using methods that worked for all of them. The trait data were either obtained from dedicated new experiments or collected from previously published studies (Lehmann et al., 2018; Zheng et al., 2018) using the set of 31 fungal strains; data origin is given in the text. With the exception of hyphal length, all traits were measured under standardized in vitro conditions which were suitable for all our fungal strains. It was not feasible to realize trait measurements in soil since it is an opaque and highly heterogeneous substrate. Instead we used potato dextrose agar, a widely used standard growth medium for fungi. By this, we ensure a consistent environmental setting for trait measurements (Aguilar-Trigueros et al., 2015; Lehmann and Rillig, 2015).

#### Morphological traits

We measured hyphal length in soil (in m g^−1^ soil); for this we used soil samples from the soil aggregate formation assay; hence we had ten replicates for each fungal strain and the control. For extracting hyphae and measuring hyphal length, 4.0 g of the experimental soil were used, and hyphae counted at 200x magnification (Tennant, 1975; Jakobsen et al., 1992). The hyphal length found in the controls was set as the background; that is, dead hyphae that were present in the soil after autoclaving.

In order to measure colony radial growth rate (in µm h^−1^), the 31 fungal strains were cultivated on full strength PDA - a rich medium generally preventing growth limitations in our fungal strains. For each fungal strain five replicates were used. For the set-up, a pre-sterilized poppy seed colonized by a fungal strain was placed in the center of a PDA plate which was then incubated for four weeks in the dark at room temperature (22°C). At day 0, 3, 5, 7, 14, 21 and 28, all plates were scanned from the back with an Epson Perfection V700 Photo Scanner (300 dpi, 16-bit, color). The pictures were analyzed in ImageJ (Schneider et al., 2012) (1.51j8) by measuring the radius in four directions (0°, 90°, 180° and 270°) with the poppy seed as center point to the colony rim. The four values were averaged. For each replicate, the mean colony radius was plotted over time to identify the linear growth phase. The slope of the linear growth phase represents the colony radial growth rate and was estimated by linear regression standardized by the length of the linear growth phase.

The data for colony biomass density (in µg mm^−2^) were obtained in an experiment in which fungal colonies were grown on PDA covered with sterilized cellophane, allowing easy extraction of fungal biomass. For each fungal strain, six replicates were set up using colonized poppy seeds, as above. When fungi reached half of their linear growth phase, colony area was measured, then biomass was harvested, dried at 45 °C and weighed. Finally, the biomass was standardized by the colony area (Reeslev and Kjoller, 1995).

Furthermore, we included data on hyphal branching angle, hyphal internodal length, hyphal diameter, mycelial complexity (box counting dimension, describing the degree of detail of a pattern), and mycelium heterogeneity (lacunarity, i.e. the gappiness or ‘rotational and translational invariance’ in a pattern (Karperien, 1999-2013)) and hyphal surface area which were collected by Lehmann et al. (2018). For further information on experimental set-up and measurements see supplementary material.

#### Chemical traits

We measured hydrophobicity of the fungal surface for fungal material using the same approach as applied for biomass density measurements, with six replicates per fungal strain. This allowed us to use medium-free fungal material. Half of an individual colony was used for the hydrophobicity test, which was done using alcohol percentage tests. This is a rapid and simple way of quantifying hydrophobicity (Chau et al., 2010). Briefly, a series of ethanol droplets (8 µl) with a concentration gradient were placed on the fungal surface to find the maximum concentration at which the droplet can retain its shape for longer than 5 seconds (Zheng et al., 2014).

Additionally, we included here the enzymatic activity data for laccase, cellobiohydrolase, leucine aminopeptidase and acid phosphatase, previously measured by Zheng et al. (Zheng et al., 2018). For further information, see supplementary material.

#### Biological trait

The palatability of the 31 fungal strains was tested in a feeding experiment with the collembolan *Folsomia candida*. We measured palatability as a proxy for assessing likely persistence of hyphae in the environment, as a way to assess possible interaction with other soil biota. Fungal mycelium was grown on glass fiber filter papers (696, VWR European Cat. No. 516-0877) cut into 1 cm² pieces of which four were placed in Petri dishes filled with plaster of Paris and charcoal (3:1 mixture). There were 31 fungal treatments and a non-fungal control (glass fiber filters only), each with eight replicates resulting in 256 experimental units. The experiment started with the addition of ten individuals of Collembola of the same age and developmental stage; the animals were previously starved for seven days. After three days of incubation in the dark at room temperature (22°C), experimental units were checked for numbers of alive Collembola and subsequently were frozen at −20°C to stop any activity. Finally, the number of fecal pellets per dish were measured and standardized by number of surviving Collembola (fecal pellets *no. of individuals^−1^).

### 2.4. Statistics

First, for investigating soil aggregate formation (SAF) capability of the 31 fungal strains, we tested fungal performances against the corresponding control samples using a generalized least square model (gls with n= 10* 32= 320) in the ‘nlme’ package (Pinheiro et al., 2018); we accounted for heteroscedasticity by implementing different variances per stratum for fungal strains by using the varIdent function (Zuur et al., 2009). To test for differences in SAF performance of different phyla we used analysis of variance (n =31) with subsequent pairwise comparisons via TukeyHSD() function. For all models, we tested for normality and homogeneity of model residuals.

Second, we applied principal components analysis to investigate the 15-dimensional trait space and the distribution of fungal strains therein. For this, we used the prcomp() function in the basic ‘stats’ package; we used z-transformed data. To reduce the dimensionality of our dataset we tested for PC axis significance via the function testdim() (Dray, 2008) in the package ’ade4’ (Chessel et al., 2004; Dray and Dufour, 2007; Dray et al., 2007). We found that the first two axes were significant and hence used these for the PCA biplot. We added species occurrence probability information to the biplot by applying the kernel density estimation following the approach of Diaz et al. (Diaz et al., 2016). For this, we used the kde() function in the package “ks” (Duong, 2018) and implemented an unconstrained bandwidth selector via the function Hpi() for our first two PC axes. We estimated contour probabilities for 0.5, 0.95 and 0.99 quantiles with the function contourLevels(). Additionally, we tested for collinearity between our 15 trait variables by using Pearson’s rho. A threshold of |rho| >0.7 was defined as an indicator of collinearity (Dormann et al., 2013).

Third, we applied a permutation-based random forest algorithm (Hapfelmeier and Ulm, 2013) to identify informative trait variables which are important for soil aggregate formation (SAF). Random forest (Breiman, 2001) is one of the machine learning algorithms with highest accuracy (Douglas et al., 2011; Crisci et al., 2012), and is capable of detecting nonlinear relationships even among higher order interactions in a nonparametric manner (Ryo and Rillig, 2017; Ryo et al., 2018), while being robust to multicollinearity (Nicodemus et al., 2010). SAF was regressed with all the trait variables, and the model performance was evaluated in terms of explanatory power (i.e. variability explained, R^2^_expl_) and predictability using out-of-bag cross validation (Breiman, 1996) (R^2^_pred_). The relative importance of the trait variables was quantified with a mean squared error measure, indicating how much each of the trait variables contributes to the model predictability (Breiman, 2001). In addition, statistical significance of each trait variable (*p* = 0.05) was tested via a permutation approach with 2000 iterations (Hapfelmeier and Ulm, 2013). The two parameters of the random forest algorithm (see(Breiman, 2001)) were tuned as follows: the number of trees in the model (ntree) was set to 100 as it made the model stable (Breiman, 2001); the number of predictors for the randomized split technique (mtry) was set to 4 (the square root of the number of predictors (Diaz-Uriarte and de Andres, 2006)).

We added the phylogeny of our 31 fungal strains as a numeric predictor variable to the random forest analysis. To do this, we calculated phylogenetic pairwise distances and fed these into PCoA via the cmdscale() function in the ‘stats’ package. We calculated the cumulative sum of the proportion of variance explained by PCo axes based on the eigenvalues and extracted the first five axes, together explaining up to 80% of phylogenetic variance (Diniz-Filho et al., 1998). After identifying the most relevant predictors, we used partial dependence plots to visualize the response-predictor relationships obtained from the random forest procedure (Hastie et al., 2009). For this, we used the plotPartialDependence() function of the package ‘mlr’ (Bischl et al., 2016).

Fourth, we tested for phylogenetic signals in our 15 trait variables (Table S3) using Moran’s I statistic -a measure of phylogenetic autocorrelation, implemented in the package ‘phylosignal’ (Keck et al., 2016).

Fifth, we ran linear regressions on SAF and the three most important predictors identified by the random forest approach and further evaluated the relationships by quantile regression with the package ‘quantreg’ (https://github.com/cran/quantreg). Analyzing response-predictor relationships at their maxima rather than at their means allows for more meaningful inferences especially for wedge-shaped data distributions (Cade et al., 1999; Cade and Noon, 2003); in these cases, unmeasured limiting factors could obscure underlying patterns. Model residuals were tested for homogeneity and normal distribution. If necessary, data were log-transformed. Sixth, we visually explored soil aggregate formation strategies exemplified by the four best and poorest performing strains via radar charts applying the eponymous function in the package ’fmsb’ (Nakazawa, 2018).

We conducted all analyses in R (R Development Core Team, 2014) (v. 3.4.1) and generated plots, if not stated otherwise, with the graphic package ‘ggplot2’ (Wickham, 2009).

## 3. Results and Discussion

### 3.1. Soil aggregate formation

We here measured soil aggregate formation (SAF) capability on a broad set of fungal strains comprising the phyla Ascomycota, Basidiomycota and Mucoromycota, revealing an overall significantly positive effect of fungi on soil aggregation: the saprobic fungi increased SAF of the tested sandy soil by 79% (confidence interval: 61 to 99%; Fig S1) compared to the non-inoculated controls. The control samples reached a SAF of 3.5% (standard deviation: 0.6) while, for the fungal treatments, we found a spectrum of SAF with means ranging from 3.7 to 10.3% with the Mucoromycota strain *Umbelopsis isabellina* and the Ascomycota strain *Cadophora sp*. at the lower and upper end, respectively (Fig. 2A). Only two strains, namely *Umbelopsis isabellina* and *Mortierella sp*.3, had a SAF performance not significantly different from the non-inoculated controls.

**Fig. 2.**
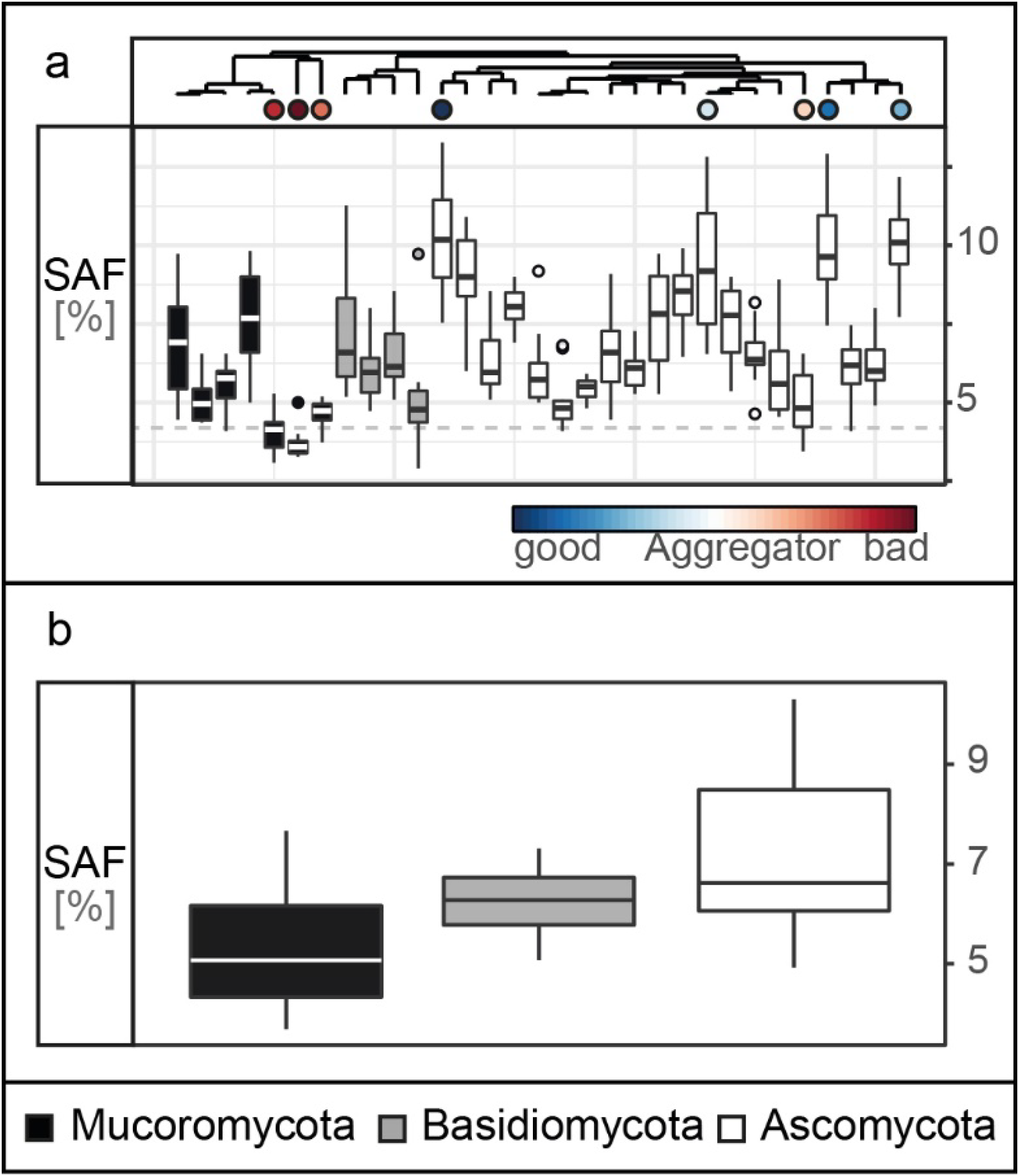
Soil aggregate formation capability. (a) Tukey boxplots of the soil aggregate formation (SAF with n= 10 *31 in %) capability of the 31 fungal strains. The dashed line represents the average SAF of the controls (n=10, mean= 3.5, SD=0.59) (b) Soil aggregate formation capability depicted on phylum level (pairwise comparisons: Ascomycota - Basidiomycota: p= 0.47, Ascomycota - Mucoromycota: p= 0.03, Basidiomycota - Mucoromycota: p= 0.66; n=31).

Our results support the general finding that filamentous soil fungi improve soil aggregation, as was shown in experiments (Martin and Anderson, 1943; Gilmour et al., 1948; Martin et al., 1958; Zheng et al., 2014) and a global data synthesis (Lehmann et al., 2017b). However, here we used for the first time a set of 31 fungal strains comprising three major fungal phyla which were all isolated from the same soil and tested in their home soil. This set was screened using a method suitable for the large number of target species. Additionally, we used a straightforward assay for testing specifically a soil aggregation process component - namely aggregate formation.

Our choice of methods also has limitations. Using this approach, we only focused on one *a apriori* size limit for newly formed aggregates, thus not capturing any dynamics in smaller sizes classes. Furthermore, the small amount of soil used in our design did not allow us to measure aggregate size distributions. As discussed previously (Aguilar-Trigueros et al., 2015). We here evaluated fungal contribution to soil aggregation in isolation, not taking into account how such effects might be modified by other soil organisms. However, such species interactions can be clearly important; for example, a recent meta-analysis revealed that soil biota combinations (e.g. bacteria-fungi mixtures) result in significantly increased soil aggregation (Lehmann et al., 2017b). Hence future studies should also consider species combinations when evaluating soil biota contributions to soil aggregation.

In our experiment, each of the three tested fungal phyla contained strains that were effective and poorly performing; however, overall, the four most efficient aggregate formers were members of the Ascomycota while three of the poorest aggregate formers belonged to the Mucoromycota (Fig. 2B). For our tested suite of fungi, we found that the Ascomycota, in general, had significantly higher SAF than the Mucoromycota. These findings correspond with previous reports (Lynch and Elliott, 1983; Tisdall et al., 2012; Lehmann et al., 2017b) and suggest that phylogeny is a strong factor determining SAF capability. However, it still remains unclear which fungal traits contribute to these phylum-specific differences and overall variability in SAF capability. Thus, in the next step, we used a trait database comprising morphological, chemical and biotic traits to explore their importance for SAF.

### 3.2. Trait collection

We included 15 fungal traits (measured on the level of a fungal individual or ‘colony’) and found strong variability in their expression across the 31 fungal strains (Fig. 3). In terms of morphological features, we found in our experiments that the measured branching angles ranged from 26 to 86° for Mucoromycota with widest and Basidiomycota with narrowest angles, while for hyphal diameter, the highest and lowest values (2.7 to 6.5 µm) were both found in the Mucoromycota. Basidiomycota had the highest internodal length (453 µm) while in Mucoromycota distances as short as 40 µm between two branches were detected. The mycelium complexity measurements revealed trait values between 1.2 (Basidiomycota) and 1.6 (Mucoromycota), where a value of 1 represents a single, unbranched hypha and a value of 2 a complex, space-filling structure. Mycelium heterogeneity varied between 0.4 and 0.7 for Basidio- and Ascomycota, respectively, with higher values indicating increasing structural gappiness. For hyphal length in soil, we found 7 to 20 m hyphae per g soil for Ascomycota and Basidiomycota, respectively, with 4.6 m g^−1^ of hyphal background. The largest hyphal surface area was found in Mucoromycota with 3.4 µm² while the smallest was detected for an Ascomycota strain with 0.8 µm². For biomass density, values ranged between 0.02 and 0.2 mg cm^−2^ for Basidiomycota and Ascomycota, respectively. Among the Mucoromycota the strain with the highest colony radial extension rate with 373 µm h^−1^ was found while the slowest extending strain was a member of the Ascomycota.

**Fig. 3.**
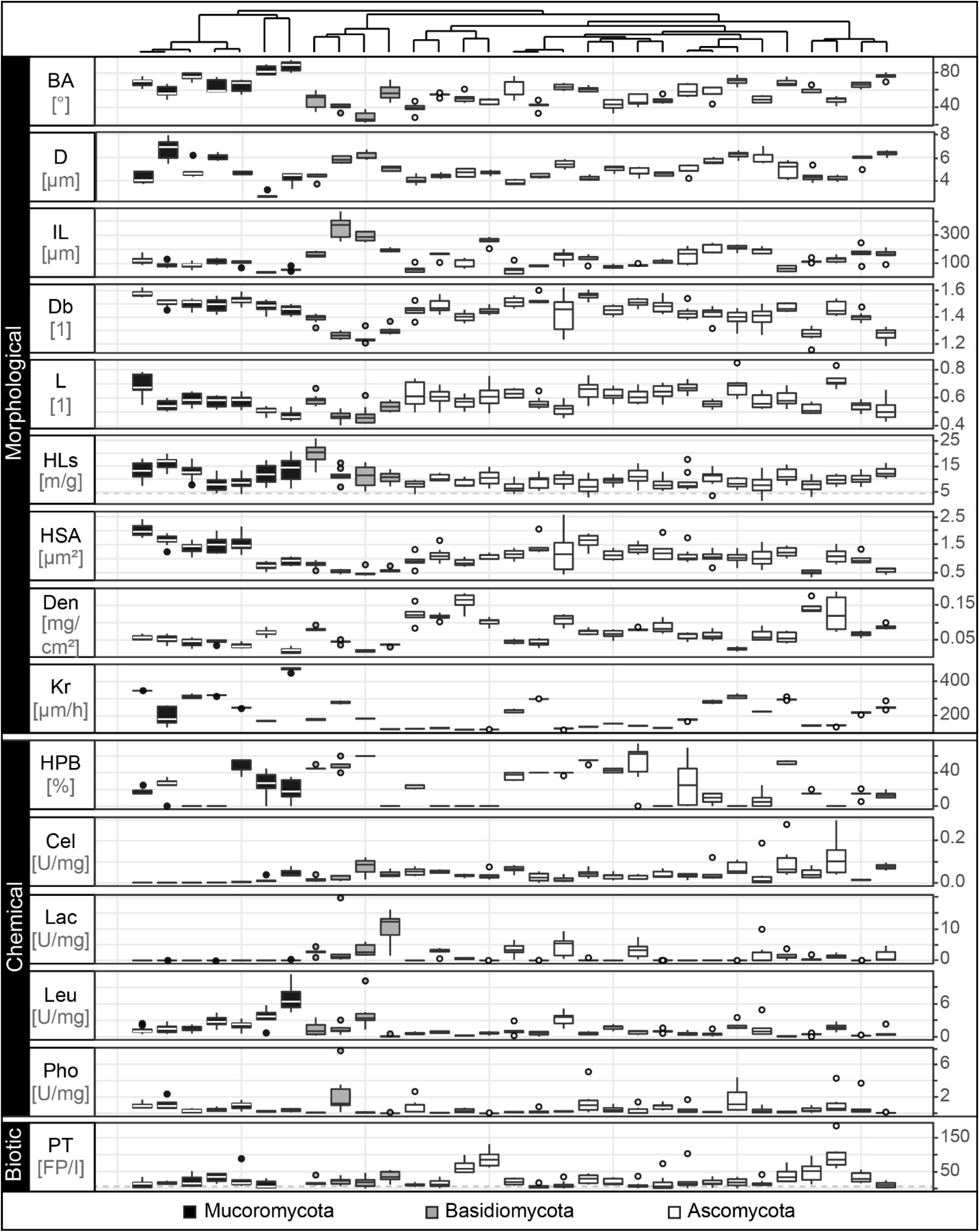
Trait distributions. Tukey boxplots of the 15 trait variables comprising morphological, chemical and biotic fungal features. Here, we present data on branching angle (BA with n= 5 in °), hyphal diameter (D with n= 5 in µm), internodal length (IL with n= 5 in µm), boxcounting dimension (Db with n= 8, unitless), lacunarity (L with n= 8, unitless), hyphal length in soil (HLs with n= 10 in m/g), hyphal surface area (HSA with n= 8 in µm²), biomass density (Den with n= 6 in mg*cm^−^²), radial colony extension rate (Kr with n= 5 in µm*h^−1^), hydrophobicity of fungal surfaces (HPB with n= 6 in % of ethanol molarity), cellobiohydrolase (Cel), laccase (Lac), leucine aminopeptidase (Leu) and acid phosphatase (Pho) activity (each with n= 5 in unit* g^−1^ dry mass) and palatability (PT with n= 8 in no. of fecal pellets per collembolan individual). The boxplots represent 25th and 75th percentile, median and outlying points. Information about phylum affiliation is colour-coded (black: Mucoromycota, grey: Basidiomycota, white: Ascomycota). The grey dashed line for the trait hyphal length in soil represents mean of corresponding trait controls.

Next, the exploration of the chemical traits revealed that across all phyla, hydrophilic mycelia could be found while Basidiomycota showed the strongest detectable mycelial hydrophobicity (60% ethanol molarity). The enzyme profiling revealed that cellobiohydrolase was not produced by Mortierellales, an order of the Mucoromycota, while the highest activity was found in the Ascomycota (0.13 U mg-1). In contrast, laccase and acid phosphatase activities were lowest in Ascomycota and highest in Basidiomycota (laccase: 0.01 to 10.4 U mg-1; acid phosphatase: 0.01 to 1.8 U mg-1). The production of leucine aminopeptidase was highest in Mucoromycota and lowest in Ascomycota (0.09 to 7.1 U mg-1).

We measured palatability as a biotic trait and found that the most and least attractive strains belonged to the Ascomycota (5 to 123 fecal pellets per individual collembolan).

Here, we established a collection of soft traits measured under standardized conditions with reproducible methods which are applicable for a broad range of fungal strains with high intra- and interspecific variability in morphological, chemical and biotic features. Our values are within the range of previously reported fungal traits (e.g. Trinci, 1969; Ho, 1978; Obert et al., 1990; Baldrian et al., 2011; Eichlerova et al., 2015).

However, it is important to note that these findings result from trait data measured on a homogenous, standardized growth substrate not accounting for the heterogeneous nature of soil with its inherent structure and also physical, chemical and biotic factors influencing the fungal trait expression. It is well known that fungal mycelia are versatile, dynamic and modular constructs; they not only modify their environment during foraging but also react to it (Ritz and Young, 2004). As demonstrated using the model organism *Rhizoctonia solani*, nutrient distribution and soil bulk density can alter e.g. hyphal growth patterns and thus mycelium density (Harris et al., 2003; Boswell et al., 2007). Future studies would need to take into account the soil heterogeneity.

### 3.3. Fungal trait space

We investigated the resulting 15-dimensional trait space and the fungal strain probability occurrence therein (Fig. 4A). We constructed the trait space by ordination (principal components analysis) and hence converted individual strains into unique trait combinations whose coordinates are determined by their trait expression (Crowther et al., 2014; Aguilar-Trigueros et al., 2015). We found that 42% of the variability in the fungal traits was accounted for in the first two PC axes which were the only significant axes (Table S4). Due to indication of strong trait correlations, we tested our data for collinearity. We detected only one case of collinearity (|pearson’s rho|>0.7) for mycelium complexity and hyphal surface area (Fig. S2).

**Fig. 4.**
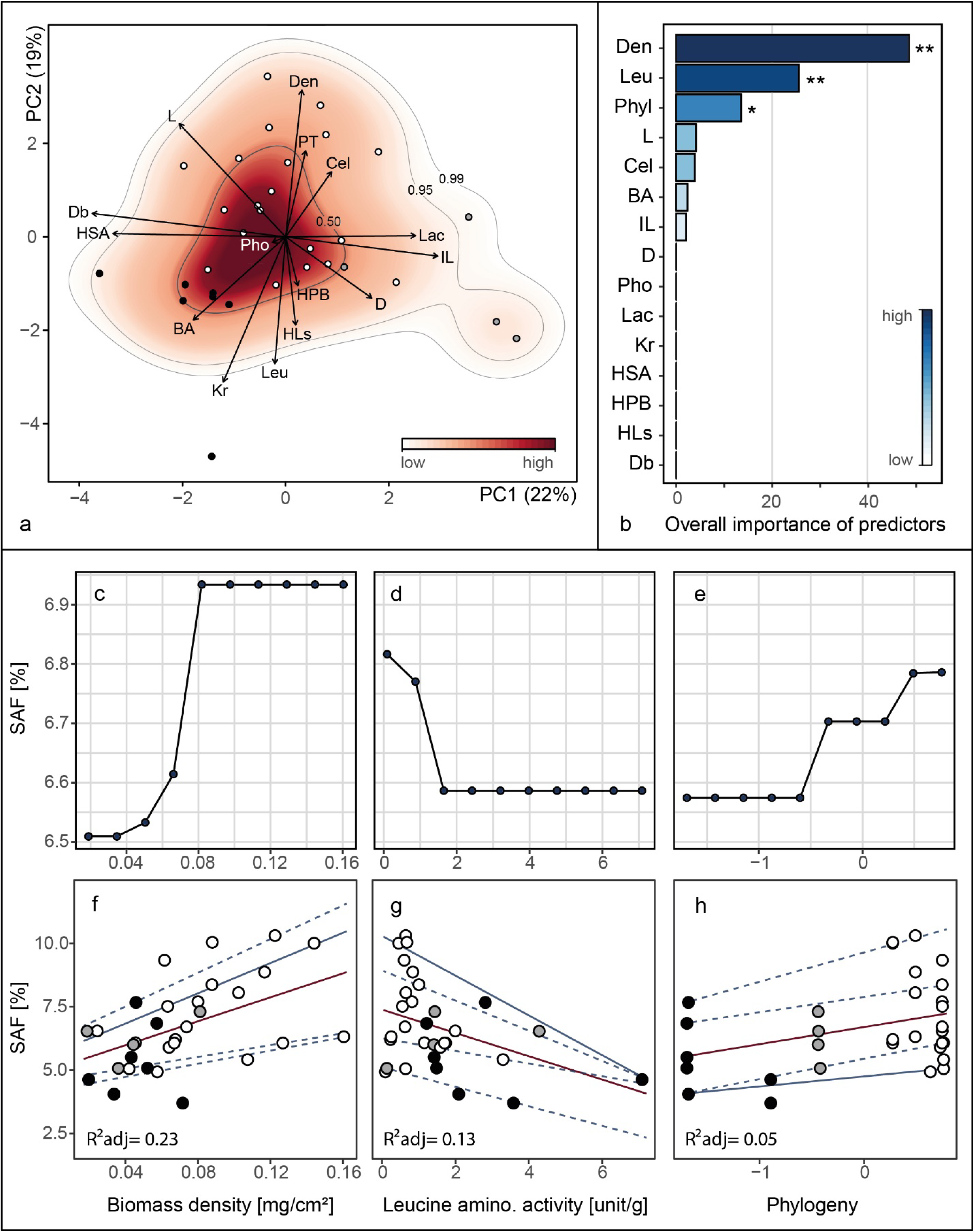
Outcomes of principal components analysis, random forest analysis and relationships between soil aggregate formation (SAF) and important trait variables. Analyses were conducted on trait mean data (n= 31). **(A)** Projection of the ordinated 31 fungal strains onto 15 trait variables comprising morphological, chemical and biotic characteristics into two dimensional trait space represented by principal component axis 1 and 2 (explaining 23 and 19% of variance, respectively). The trait variables are branching angle (BA), hyphal diameter (D), internodal length (IL), boxcounting dimension (Db), lacunarity (L), hyphal length in soil (HLs), hyphal surface area (HSA), biomass density (Den), radial colony extension rate (Kr), hydrophobicity of fungal surfaces (HPB), cellobiohydrolase (Cel), laccase (Lac), leucine aminopeptidase (Leu) and acid phosphatase (Pho) activity and palatability (PT). Arrows indicate direction and weight of trait vectors. Colour gradient represents probability of species occurrence (white = low, red = high) in the trait space, with the contour lines denoting the 0.50, 0.95 and 0.99 quantiles of kernel density estimation (see materials and methods section). **(B)** Overall importance of trait variables for soil aggregate formation capability with R^2^_expl_ = 0.36, R^2^_pred_ = 0.13 and three statistically significant predictor variables. Asterisks denote significance level: *** < 0.0001, ** 0.001, * 0.01,. 0.5. Pairwise phylogenetic distance was included as PCo-axes (see materials and methods section). **(C-E)** Partial dependence plots for the three most important and significant trait variables identified by random forest approach. The x-axis labels are identical with panels F, G and H, respectively. **(F-H)** Relationships between SAF and the three most important trait variables. Corresponding regression statistics can be found in Table S5. Phylum affiliation of fungal strains is colour-coded (black: Mucoromycota, grey: Basidiomycota, white: Ascomycota). Red and blue lines represent linear and quantile regression lines, respectively. The line type depicts significance of regression lines with solid < 0.05 and dashed > 0.05.

Evaluating the species occurrence, we found that Ascomycota strains were distributed in the lower half of the PC plane whereas the Mucoromycota were localized in the upper left quadrant mainly characterized by hyphal branching angle, colony radial growth rate and leucine aminopeptidase activity. In the upper right quadrant, the Basidiomycota grouped driven by hyphal internodial length and lacunarity. There was a clear separation of the phyla detectable for PC axis 1 with Ascomycota flanked by Mucoromycota and Basidiomycota but only a marginal separation between Ascomycota and Mucoromycota on PC axis 2 (Fig. S3). In general, the trait space revealed a high versatility in our fungal set with no clear syndromes. However, on the phylum level a clear separation between the three phyla was evident (Fig. S3). In the next step, we investigated the importance of the collected fungal traits on SAF using the random forest approach. Considering the strong impact of phylum on SAF and phylogenetic separation in the trait space, we included phylogenetic pairwise distances as an additional variable (potentially also capturing not explicitly measured variables) in the following analyses.

### 3.4. Fungal trait contributions to soil aggregate formation

The random forest algorithm (explanatory power: 36% and predictability: 13%), identified three significant trait variables: colony biomass density, leucine aminopeptidase activity and phylogeny (relative importance: 48%, 25% and 13%, explanatory power of each: 17.3%, 9%, 4.7%; Fig. 4B).

To visualize the modeled relationship between SAF and the important variables we used partial dependence plots. After taking into account the effects of all predictors except for the variable of interest (colony biomass density, leucine aminopeptidase activity or phylogeny, respectively), partial dependence plots depict the relationships between the predictor and the response variable (SAF). We found that SAF increased with increasing colony biomass density (Fig. 4C) but decreased with increasing leucine aminopeptidase activity (Fig. 4D). Across the phylogeny, from Mucoromycota to Ascomycota, we found a positive relationship with SAF (Fig. 4E). These findings were supported by linear and quantile regression analyses (Fig. 4F to 4H, Table S5). Here, we found that the relationship between SAF and colony biomass density was best represented by mean regression. For the relationships between SAF and leucine aminopeptidase activity as well as SAF and phylogeny, the 0.95 and 0.05 quantile, respectively, showed the highest fit.

Our analyses revealed that fungal strains belonging to the Ascomycota that have high biomass density and low leucine aminopeptidase activity have the highest probability to form aggregates compared to other strains. Furthermore, we found that a colony biomass density above 0.08 mg cm^−2^ and a leucine aminopeptidase activity less than 1.8 U g^−1^ do not further improve SAF (Fig. 4C and 4D).

Our findings further support the assumption that phylogeny influences aggregate forming capability of fungi (Fig. 3B and Fig. 4H). We interpret this to mean that traits (including unmeasured traits) expressed by strains of this phylum contribute to this beneficial impact on soil aggregation. Considering all possible traits and their expression, the four most efficient aggregate former were all Ascomycota with low leucine aminopeptidase activity and dense mycelia.

A densely growing fungus likely can more intensively cross-link and enmesh particles with its hyphae, and thus perhaps is more effective at contributing to the formation of macroaggregates; however, so far there has not been direct evidence of this. Interestingly, the total amount of hyphae produced was not an important explanatory variable (Fig. 2; HLs = hyphal length in soil) suggesting that a critical local density is much more important than total hyphal production. This also explains results from previous experiments, where total hyphal length or biomass did not predict soil aggregation effects (e.g. Piotrowski et al., 2004). Fungi with high biomass density had low radial colony extension rate (Fig. S2); thus it can be expected that their positive effect on SAF is highly localized not reaching beyond their area of mycelial influence.

Fungi with low leucine aminopeptidase activity are inefficient in hydrolyzing peptides and thus degrading organic matter components, which may be functioning as glues and cementing agents in aggregates (Chenu, 1989; Caesar-TonThat and Cochran, 2000; Daynes et al., 2012). Fungi with either one of these traits are more likely able to bring soil particles and aggregates together via their hyphae; lacking the enzyme to degrade organic matter holding together aggregates also contributes to this effect.

This holds true especially in soils with high sand content as-used in our assay. In such soils, fungi are an essential factor in soil aggregation mainly via physical and chemical interactions of hyphae with sand particles forming and stabilizing the otherwise unstable substrate (Sutton and Sheppard, 1976; Forster and Nicolson, 1981). We here chose the soil from which our fungi were originally cultured. However, soil type as a major variable affecting fungi and their soil aggregation capability has to be the main target of future studies.

After identifying the most important fungal traits for SAF, we focused on those fungi that are present at the lower and upper end of the SAF spectrum. The most efficient strains were all members of the Ascomycota (*Cadophora sp*., *Pleosporales sp*., *Alternaria sp*., *Fusarium sp*.) while the group of the poor performer comprised mainly Mucoromycota but also one ascomycete (*Umbelopsis isabellina*, *Mortierella sp*. (no. 3), *Mucor fragilis*, *Truncatella angustata* (Fig.1 and Fig.3). As expected, the efficient soil aggregate forming strains had high biomass density but low leucine aminopeptidase activity (Fig. 5). The opposite was true for the poor performers. In addition to these two clear features, the efficient strains tended to have lower colony radial growth rates, hyphal surface area and surface hydrophobicity, but had larger hyphal diameters and more heterogeneously structured mycelia as the four poorest soil aggregators.

**Fig. 5.**
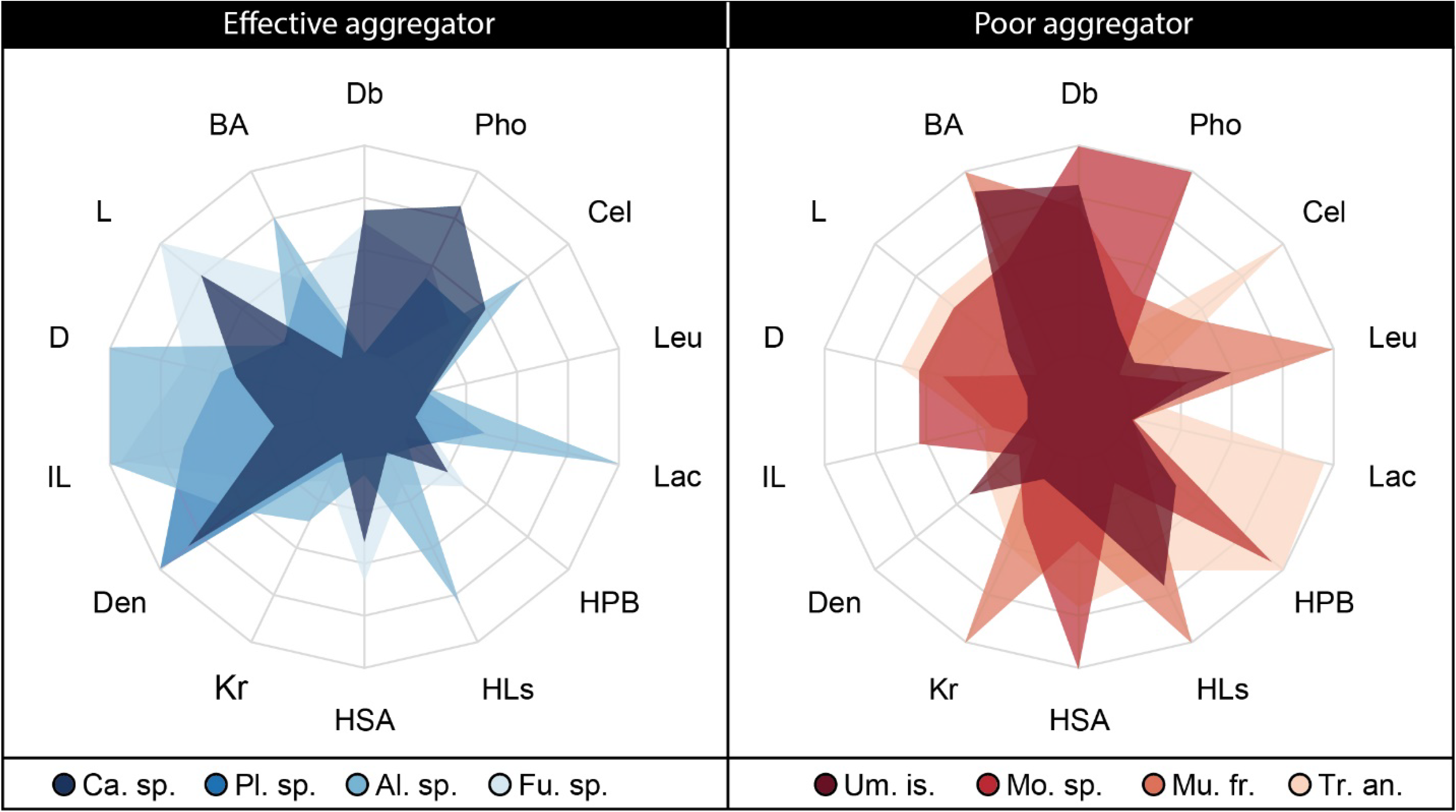
Radar plot depicting trait expressions for the four best and four poorest soil aggregate forming fungal strains.

## 4. Conclusions

Our results yield new insights into fungal traits important for soil aggregation, and thus also shed light on mechanisms of soil aggregation. Clearly, future work should focus on hyphal density as a key trait. In an applied context of restoration and agriculture, our trait information can be incorporated in management practices affecting the fungal environment in soil to favor the development of more dense fungal mycelia by e.g. carbon input or through a screen for isolates exhibiting desired traits under the soil conditions in which they will be used.

Even though we here focused on saprobic soil fungi, some aspects may also be generalizable to other fungal groups. For example, future work should test if hyphal density is also a better predictor for soil aggregation ability than hyphal biomass production in arbuscular mycorrhizal fungi. On the other hand, it will also be important to extend the dataset of fungal traits and soil aggregation beyond soil saprobes, since the relative importance of traits and trait combinations could vary; for example, since arbuscular mycorrhizal fungi have limited enzymatic abilities (Tisserant et al., 2013), this trait would play no role in that particular group. In the end, our study demonstrates the power of employing a trait-based approach to tackle biological mechanisms of soil aggregation; this can now also be extended to organism groups other than fungi.

## Acknowledgements

This work was supported by the Deutsche Forschungsgemeinschaft (RI 1815/16-1).

## Author contributions

A.L. designed and performed the research; W.Z. and M.R. contributed analytical tools; A.L, W.Z., K.S, R.R. and S.M. provided experimental data; J.R. created the phylogenetic tree; A.L. and M.C.R. wrote the manuscript; all authors contributed to the final version of the manuscript.

## Additional information

Supplementary information is available for this paper (Supplementary Information and Supplementary Data).

## References

Aguilar-Trigueros, C.A., Hempel, S., Powell, J.R., Anderson, I.C., Antonovics, J., Bergmann, J., Cavagnaro, T.R., Chen, B.D., Hart, M.M., Klironomos, J., Petermann, J.S., Verbruggen, E., Veresoglou, S.D., Rillig, M.C., 2015. Branching out: Towards a trait-based understanding of fungal ecology. Fungal Biology Reviews 29, 34–41.

Andrade-Linares, D.R., Veresoglou, S.D., Rillig, M.C., 2016. Temperature priming and memory in soil filamentous fungi. Fungal Ecology 21, 10–15.

Baldrian, P., Voriskova, J., Dobiasova, P., Merhautova, V., Lisa, L., Valaskova, V., 2011. Production of extracellular enzymes and degradation of biopolymers by saprotrophic microfungi from the upper layers of forest soil. Plant and Soil 338, 111–125.

Bardgett, R.D., van der Putten, W.H., 2014. Belowground biodiversity and ecosystem functioning. Nature 515, 505–511.

Bischl, B., Lang, M., Kotthoff, L., Schiffner, J., Richter, J., Studerus, E., Casalicchio, G., Jones, Z.M., 2016. mlr: Machine Learning in R. Journal of Machine Learning Research 17.

Boswell, G.P., Jacobs, H., Ritz, K., Gadd, G.M., Davidson, F.A., 2007. The development of fungal networks in complex environments. Bulletin of Mathematical Biology 69, 605–634.

Breiman, L., 1996. Out-of-bag estimation. https://www.stat.berkeley.edu/users/breiman/OOBestimation.pdf.

Breiman, L., 2001. Random Forests. Machine Learning 45, 5–32.

Bronick, C.J., Lal, R., 2005. Soil structure and management: a review. Geoderma 124, 3–22.

Cade, B.S., Noon, B.R., 2003. A gentle introduction to quantile regression for ecologists. Frontiers in Ecology and the Environment 1, 412–420.

Cade, B.S., Terrell, J.W., Schroeder, R.L., 1999. Estimating effects of limiting factors with regression quantiles. Ecology 80, 311–323.

Caesar-TonThat, T.C., Cochran, V.L., 2000. Soil aggregate stabilization by a saprophytic lignin- decomposing basidiomycete fungus - I. Microbiological aspects. Biology and Fertility of Soils 32, 374–380.

Chau, H.W., Goh, Y.K., Si, B.C., Vujanovic, V., 2010. Assessment of alcohol percentage test for fungal surface hydrophobicity measurement. Letters in Applied Microbiology 50, 295–300.

Chenu, C., 1989. Influence of a fungal polysaccharide, scleroglucan, on clay microstructure. Soil Biology & Biochemistry 21, 299–305.

Chessel, D., Dufour, A.B., Thioulouse, J., 2004. The ade4 package - I: One-table methods. R News 4, 5–10.

Crisci, C., Ghattas, B., Perera, G., 2012. A review of supervised machine learning algorithms and their applications to ecological data. Ecological Modelling 240, 113–122.

Crowther, T.W., Maynard, D.S., Crowther, T.R., Peccia, J., Smith, J.R., Bradford, M.A., 2014. Untangling the fungal niche: the trait-based approach. Frontiers in Microbiology 5, 1–12.

Daynes, C.N., Zhang, N., Saleeba, J.A., McGee, P.A., 2012. Soil aggregates formed in vitro by saprotrophic Trichocomaceae have transient water-stability. Soil Biology & Biochemistry 48, 151–161.

Diaz-Uriarte, R., de Andres, S.A., 2006. Gene selection and classification of microarray data using random forest. BMC Bioinformatics 7.

Diaz, S., Kattge, J., Cornelissen, J.H.C., Wright, I.J., Lavorel, S., Dray, S., Reu, B., Kleyer, M., Wirth, C., Prentice, I.C., Garnier, E., Bonisch, G., Westoby, M., Poorter, H., Reich, P.B., Moles, A.T., Dickie, J., Gillison, A.N., Zanne, A.E., Chave, J., Wright, S.J., Sheremet’ev, S.N., Jactel, H., Baraloto, C., Cerabolini, B., Pierce, S., Shipley, B., Kirkup, D., Casanoves, F., Joswig, J.S., Gunther, A., Falczuk, V., Ruger, N., Mahecha, M.D., Gorne, L.D., 2016. The global spectrum of plant form and function. Nature 529, 167–173.

Diniz-Filho, J.A.F., de Sant’Ana, C.E.R., Bini, L.M., 1998. An eigenvector method for estimating phylogenetic inertia Evolution 52, 1247–1262.

Dormann, C.F., Elith, J., Bacher, S., Buchmann, C., Carl, G., Carre, G., Marquez, J.R.G., Gruber, B., Lafourcade, B., Leitao, P.J., Munkemuller, T., McClean, C., Osborne, P.E., Reineking, B., Schroder, B., Skidmore, A.K., Zurell, D., Lautenbach, S., 2013. Collinearity: a review of methods to deal with it and a simulation study evaluating their performance. Ecography 36, 27–46.

Douglas, P.K., Harris, S., Yuille, A., Cohen, M.S., 2011. Performance comparison of machine learning algorithms and number of independent components used in fMRI decoding of belief vs. disbelief. NeuroImage 56, 544–553.

Dray, S., 2008. On the number of principal components: A test of dimensionality based on measurements of similarity between matrices. Computational Statistics & Data Analysis 52, 2228–2237.

Dray, S., Dufour, A.B., 2007. The ade4 package: Implementing the duality diagram for ecologists. Journal of Statistical Software 22, 1–20.

Dray, S., Dufour, A.B., Chessel, D., 2007. The ade4 package-II: Two-table and K-table methods. R News 7, 47–52.

Drummond, A.J., Rambaut, A., 2007. BEAST: Bayesian evolutionary analysis by sampling trees. BMC Evolutionary Biology 7.

Duong, T., 2018. ks: Kernel smoothing, R package version 1.11.0. ed.

Edgar, R.C., 2004. MUSCLE: multiple sequence alignment with high accuracy and high throughput. Nucleic Acids Research 32, 1792–1797.

Eichlerova, I., Homolka, L., Zifcakova, L., Lisa, L., Dobiasova, P., Baldrian, P., 2015. Enzymatic systems involved in decomposition reflects the ecology and taxonomy of saprotrophic fungi. Fungal Ecology 13, 10–22.

Forster, S.M., Nicolson, T.H., 1981. Microbial aggregation of sand in a maritime dune succession. Soil Biology and Biochemistry 13, 205–208.

Gams, W., Domsch, K.H., 1967. Beitrage zur Anwendung der Bodenwaschtechnik für die Isolierung von Bodenpilzen. Arch. Mikrobiol. 58, 134–144.

Gilmour, C.M., Allen, O.N., Truog, E., 1948. Soil aggregation as influenced by the growth of mold species, kind of soil, and organic matter. Soil Science Society of America Proceedings 13, 292–296.

Hapfelmeier, A., Ulm, K., 2013. A new variable selection approach using Random Forests. Computational Statistics & Data Analysis 60, 50–69.

Harris, K., Young, I.M., Gilligan, C.A., Otten, W., Ritz, K., 2003. Effect of bulk density on the spatial organisation of the fungus Rhizoctonia solani in soil. FEMS Microbiology Ecology 44, 45–56.

Hastie, T., Tibshirani, R., Friedman, J., 2009. The elements of statistical learning: Data mining, inference, and prediction, 2 ed. Springer, New York.

Ho, H.H., 1978. Hyphal branching systems in Phytophthora and other Phcomycetes. Mycopathologia 64, 83–86.

Jakobsen, I., Abbott, L.K., Robson, A.D., 1992. External hyphae of vesicular-arbuscular mycorrhizal fungi associated with *Trifolium subterraneum* L. 1. Spread of hyphae and phosphorus inflow into roots. New Phytologist 120, 371–380.

Karperien, A., 1999–2013. FracLac for ImageJ.

Keck, F., Rimet, F., Bouchez, A., Franc, A., 2016. phylosignal: an R package to measure, test, and explore the phylogenetic signal. Ecology and Evolution 6, 2774–2780.

Lehmann, A., Leifheit, E.F., Rillig, M.C., 2017a. Mycorrhizas and soil aggregation, In: Johnson, N.C., Gehring, C., Jansa, J. (Eds.), Mycorrhizal mediation of soil, 1 ed. Elsevier, Amsterdam, Netherlands.

Lehmann, A., Rillig, M.C., 2015. Understanding mechanisms of soil biota involvement in soil aggregation: A way forward with saprobic fungi? Soil Biology and Biochemistry 88, 298–302.

Lehmann, A., Zheng, W., Soutschek, K., Rillig, M.C., 2018. How to build a mycelium: tradeoffs in fungal architectural traits. bioRxiv.

Lehmann, A., Zheng, W.S., Rillig, M.C., 2017b. Soil biota contributions to soil aggregation. Nature Ecology & Evolution 1, 1828–1835.

Lynch, J.M., Elliott, L.F., 1983. Aggregate stabilization of volcanic ash and soil during microbial-degradation of straw. Applied and Environmental Microbiology 45, 1398–1401.

Martin, J.P., Ervin, J.O., Shepherd, R.A., 1958. Decomposition and aggregating effect of fungus cell material in soil. Soil Science Society of America Journal.

Martin, T.L., Anderson, D.A., 1943. Organic matter decomposition, mold flora, and soil aggregation relationships. Soil Science Society of America Journal 7, 215–217.

Nakazawa, M., 2018. fmsb: Functions for Medical Statistics Book with some Demographic Data, 0.6.3 ed.

Nascimento, F.F., dos Reis, M., Yang, Z.H., 2017. A biologist’s guide to Bayesian phylogenetic analysis. Nature Ecology & Evolution 1, 1446–1454.

Nicodemus, K.K., Malley, J.D., Strobl, C., Ziegler, A., 2010. The behaviour of Random Forest permutation-based variable importance measures under predictor correlation. BMC Bioinformatics 11.

Obert, M., Pfeifer, P., Sernetz, M., 1990. Microbial growth patterns described by fractal geometry. Journal of Bacteriology 172, 1180–1185.

Pinheiro, J., Bates, D., DebRoy, S., Sarkar, D., Team, R.C., 2018. nlme: Linear and Nonlinear Mixed Effects Models, 3.1-137 ed.

Piotrowski, J.S., Denich, T., Klironomos, J.N., Graham, J.M., Rillig, M.C., 2004. The effects of arbuscular mycorrhizas on soil aggregation depend on the interaction between plant and fungal species. New Phytologist 164, 365–373.

R Development Core Team, 2014. R: A language and environment for statistical computing, 3.4.1 ed.

Reeslev, M., Kjoller, A., 1995. Comparison of biomass dry weights and radial growth-rates of fungal colonies on media solidified with different gelling compounds. Applied and Environmental Microbiology 61, 4236–4239.

Rillig, M.C., Aguilar-Trigueros, C.A., Bergmann, J., Verbruggen, E., Veresoglou, S.D., Lehmann, A., 2015. Plant root and mycorrhizal fungal traits for understanding soil aggregation. New Phytologist 205, 1385–1388.

Ritz, K., Young, I.M., 2004. Interactions between soil structure and fungi. Mycologist 18, 52–59.

Ryo, M., Harvey, E., Robinson, C.T., Altermatt, F., 2018. Nonlinear higher order abiotic interactions explain riverine biodiversity. Journal of Biogeography 45, 628–639.

Ryo, M., Rillig, M.C., 2017. Statistically reinforced machine learning for nonlinear patterns and variable interactions. Ecosphere 8.

Schneider, C.A., Rasband, W.S., Eliceiri, K.W., 2012. NIH Image to ImageJ: 25 years of image analysis. Nature Methods 9, 671–675.

Siddiky, M.R.K., Kohler, J., Cosme, M., Rillig, M.C., 2012a.) Soil biota effects on soil structure: Interactions between arbuscular mycorrhizal fungal mycelium and collembola. Soil Biology & Biochemistry 50, 33–39.

Siddiky, M.R.K., Schaller, J., Caruso, T., Rillig, M.C., 2012b. Arbuscular mycorrhizal fungi and collembola non-additively increase soil aggregation. Soil Biology & Biochemistry 47, 93–99.

Six, J., Bossuyt, H., Degryze, S., Denef, K., 2004. A history of research on the link between (micro)aggregates, soil biota, and soil organic matter dynamics. Soil & Tillage Research 79, 7–31.

Spatafora, J.W., Chang, Y., Benny, G.L., Lazarus, K., Smith, M.E., Berbee, M.L., Bonito, G., Corradi, N., Grigoriev, I., Gryganskyi, A., James, T.Y., O’Donnell, K., Roberson, R.W., Taylor, T.N., Uehling, J., Vilgalys, R., White, M.M., Stajich, J.E., 2016. A phylum-level phylogenetic classification of zygomycete fungi based on genome-scale data. Mycologia 108, 1028–1046.

Sutton, J.C., Sheppard, B.R., 1976. AGGREGATION OF SAND-DUNE SOIL BY ENDOMYCORRHIZAL FUNGI. Canadian Journal of Botany-Revue Canadienne De Botanique 54, 326–333.

Tennant, D., 1975. Test of a modified line intersect method of estimating root length. Journal of Ecology 63, 995–1001.

Thorn, R.G., Reddy, C.A., Harris, D., Paul, E.A., 1996. Isolation of saprophytic basidiomycetes from soil. Applied and Environmental Microbiology 62, 4288–4292.

Tisdall, J.M., Nelson, S.E., Wilkinson, K.G., Smith, S.E., McKenzie, B.M., 2012. Stabilisation of soil against wind erosion by six saprotrophic fungi. Soil Biology & Biochemistry 50, 134–141.

Tisdall, J.M., Oades, J.M., 1982. Organic-matter and water-stable aggregates in soils. Journal of Soil Science 33, 141–163.

Tisserant, E., Malbreil, M., Kuo, A., Kohler, A., Symeonidi, A., Balestrini, R., Charron, P., Duensing, N., Frei dit Frey, N., Gianinazzi-Pearson, V., Gilbert, L.B., Handa, Y., Herr, J.R., Hijri, M., Koul, R., Kawaguchi, M., Krajinski, F., Lammers, P.J., Masclaux, F.G., Murat, C., Morin, E., Ndikumana, S., Pagni, M., Petitpierre, D., Requena, N., Rosikiewicz, P., Riley, R., Saito, K., San Clemente, H., Shapiro, H., van Tuinen, D., Bécard, G., Bonfante, P., Paszkowski, U., Shachar-Hill, Y.Y., Tuskan, G.A., Young, J.P.W., Sanders, I.R., Henrissat, B., Rensing, S.A., Grigoriev, I.V., Corradi, N., Roux, C., Martin, F., 2013. Genome of an arbuscular mycorrhizal fungus provides insight into the oldest plant symbiosis. Proceedings of the National Academy of Sciences 110, 20117–20122.

Trinci, A.P.J., 1969. A kinetic study of the growth of *Aspergillus nidulans* and other fungi. Journal of Genetic Microbiology 57, 11–24.

Wall, D.H., Nielsen, U.N., Six, J., 2015. Soil biodiversity and human health. Nature 528, 69–76.

Wickham, H., 2009. ggplot2: Elegant graphics for data analysis. Springer, New York.

Young, I.M., Crawford, J.W., 2004. Interactions and self-organization in the soil-microbe complex. Science 304, 1634–1637.

Zheng, W., Lehmann, A., Ryo, M., Valyi, K., Rillig, M.C., 2018. Growth rate trades off with enzymatic investment in soil filamentous fungi. bioRxiv.

Zheng, W.S., Morris, E.K., Lehmann, A., Rillig, M.C., 2016. Interplay of soil water repellency, soil aggregation and organic carbon. A meta-analysis. Geoderma 283, 39–47.

Zheng, W.S., Morris, E.K., Rillig, M.C., 2014. Ectomycorrhizal fungi in association with Pinus sylvestris seedlings promote soil aggregation and soil water repellency. Soil Biology & Biochemistry 78, 326–331.

Zuur, A.F., Ieno, E.N., Walker, N.J., Saveliev, A.A., Smith, G.M., 2009. Mixed effects models and extensions in ecology with R. Springer, UK, pp. 75–76.

